# Optimization of selective phenotyping and population design for genomic prediction

**DOI:** 10.1101/172064

**Authors:** Nicolas Heslot, Vitaliy Feoktistov

## Abstract

Calibration population design for genomic prediction has attracted a lot of interest in the plant and animal breeding literature. In this article we present an efficient optimization method to select a subset of preexisting individuals to phenotype. Application to the choice of maize hybrids to create and phenotype, to best predict the unobserved hybrid combination, is demonstrated using real data and simulations. Further, the proposed method is extended to optimize the choice of a connected population design before crosses are actually made. Population design is optimized to maximize efficiency of recurrent selection with genomic prediction. Validation results using real data and simulations are presented.

## I. INTRODUCTION

Genomic prediction [1], the use of high density whole genome markers to predict performance of both observed and unobserved individuals has fostered great interest in the plant breeding community [2]–[5]. A critical feature of any genomic prediction strategy is the population design [6], [7]. Several situations may arise. Marker data might be available on a large number of individuals but budget or logistical constraints prevent phenotyping of the whole population [8]. Methods for optimal selection of individuals to phenotype to best predict the unobserved is then needed. Similarly, in hybrid breeding, male and female inbreds are usually available with high density markers. In most cases, it is unfeasible to field test all of the possible hybrid combination. Molecular markers can be used here as well to predict unobserved hybrid combinations, often referred as hybrid prediction [9]. It is of interest to identify which hybrids to produce and phenotype to best predict the unobserved ones.

Another population design situation occurs in marker-assisted recurrent selection breeding schemes. Those schemes use markers to select unphenotyped individuals and then crossing them to generate a new generation of selection candidates. Such schemes initially used bi-parental or multipopulation QTL (quantitative trait loci) detection and then attempted to pyramid QTLs [10]. A few validation experiments of marker-assisted recurrent selection with genomic prediction are already published [11]–[14]. Identifying reliable training population design is critical for the efficiency of those schemes. Empirical results [7], [13], [15], [16] show highly variable prediction accuracies within families. Simulations results [17] also show highly variable accuracies when calibration populations are small and disconnected. Solutions are not straightforward because the only available data is high density markers on potential founders of the population. This problem was studied extensively in the context of multi-parental QTL (quantitative trait loci) mapping [18].

## II. MATERIALS AND METHODS

### A. Optimization of selective phenotyping

Mixed model based criteria [19] were proposed to optimize the choice of individuals to phenotype using as an input the expected trait heritability and markers or pedigree data [20]–[22]. Such optimization is a part of a broader class of so called “model-based design” also used to optimize field trials design [23]–[25].

For the mixed model

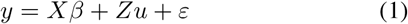

where *y* – is a vector of phenotype, *X* – a design matrix for the fixed effects, *β*, *Z* – a design matrix for the random genetic effect *u*, so that *u* is distributed multivariate normal with covariance *G*, and *ε* – the error term is multivariate normal with covariance
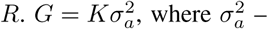
is additive genetic variance and *K* is kinship which can be based on pedigree (referred as the relationship matrix) or on markers (then often called the realized relationship matrix). If *W* is the centered markers score matrix with as many rows as individuals and as many columns as markers then

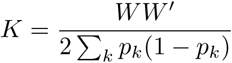

where *p_k_* – is the frequency of the minor allele [26].

From this model (Eq. 1), the mixed model equations can be written:

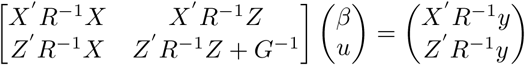

In a general case, the prediction error variance PEV for the genetic effect is:

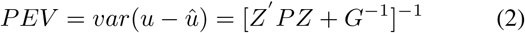

with *P* = *R*^−1^ – *R*^−1^ *X*(*X′R*^−1^*X*)^−1^*X′R*^−1^ [27]. That formulation can be re-derived directly from the mixed model equation using the matrix inversion lemma. No normality hypothesis or else are needed.

By minimizing the trace of the *PEV* (prediction error variance) matrix, or other metrics derived from that matrix, it is possible to optimize the experimental design for the precision of prediction of the genetic effects. Such metrics will be called *objective function* in the rest of the paper. Different designs imply changes in *X* and *Z*.

This formulation was used initially to optimize choice of pre-existing individuals to phenotype [20], [21]. However, evaluation of any given design is computationally intensive. For optimization speed of solution evaluation is critical. Akdemir in [28] demonstrated that the objective function above (Eq. 2) could be equivalently formulated in a more computationally efficient form. To do so, the model (Eq. 1) was rewritten at the marker effect level rather than at the individual level, with the assumptions than *X* is only an intercept and the errors are iid. The mixed model is now

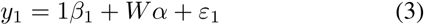

with *y*_1_ vector of phenotype, *β*_1_ intercept, *W* marker design matrix, *α* marker effects and *ε*_1_ residuals normally distributed with mean 0 and variance
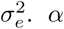
is normally distributed with mean 0 and variance
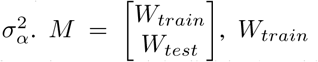
is the matrix of marker data for phenotyped individuals while *M_test_* is the matrix of marker data for unphenotyped individuals. Then the *PEV* matrix can be written *PEV^ridge^* = (1,*W_test_*)[(1,*W_train_*)′ (1, *W_train_*) + λ*I*]^−^ (1, *W_test_*)′, where ^−^ denote the pseudo inverse of a matrix and
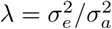
the so-called shrinkage factor. The objective function will again be the trace of the *PEV^ridge^* matrix. To speed up computation, *W* can be approximated by using the first few principal components [29]. Let *Q* the matrix of principal components of *W* partitioned as
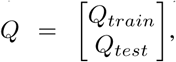
then *PEV^ridge^* ≈ (1,*Q_test_*)[(1, *Q_train_*)′ (1,*Q_train_*) + λ*I*]^−^(1, *Q_test_*)′.

This formulation is much faster to compute than the previous one, enabling more design evaluations with fixed computing resources.

An optimal solution is then searched using an optimization algorithm taking as an input λ, a total phenotyping budget and optionally constraints about individuals to be included in the solution, such as the breeder’s current most promising selection candidates or excluded from phenotyping (e.g. lack of seeds). A recent paper [30] demonstrated the use of a genetic algorithm to solve this problem.

*Optimization algorithm:* The problem is modeled as a binary optimization problem with *n* dimensional solution vector *sol* ∈ 𝓑^*n*^ ⊂ {0,1}^*n*^, where *sol_j_* = 0 means the individual belongs to the *test* set *M_test_*, and *sol_j_* = 1 individual refers to the *train* set *M_train_*, and under the constraint
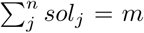
defining a phenotyping budget *m* < *n*.

Two modes of forming test set is implemented: (1) *all* the population of *n* individuals, (2) (*all* – *train*) individuals only.

Other optional constraints on individuals to be systematically phenotyped or not is handled as a fixed sub-vector of the solution vector.

To efficiently solve this problem the concept of Differential Evolution [31], one of the very promising metaheuristics, was used to create a new algorithm for binary optimization.

In order to deal with a binary search space, we invented a new operator instead of *differentiation* [32], the principal operator of Differential Evolution. We called it *activation.* An operator in Differential Evolution is a function which map one or several current solution vectors to a new vector. The newly obtained vector after a sequence of operators is usually called a trial individual.

As in any Differential Evolution (DE) algorithm, the population of *NP* individuals^1^ is randomly initialized with respect to constraints, in this case the budget constraint only. Then at each iteration, and for each of the *NP* individuals the following operators are successively executed: activation, crossover, restoration, evaluation, and selection (see Alg. 1).

#### Algorithm 1

Binary Differential Evolution

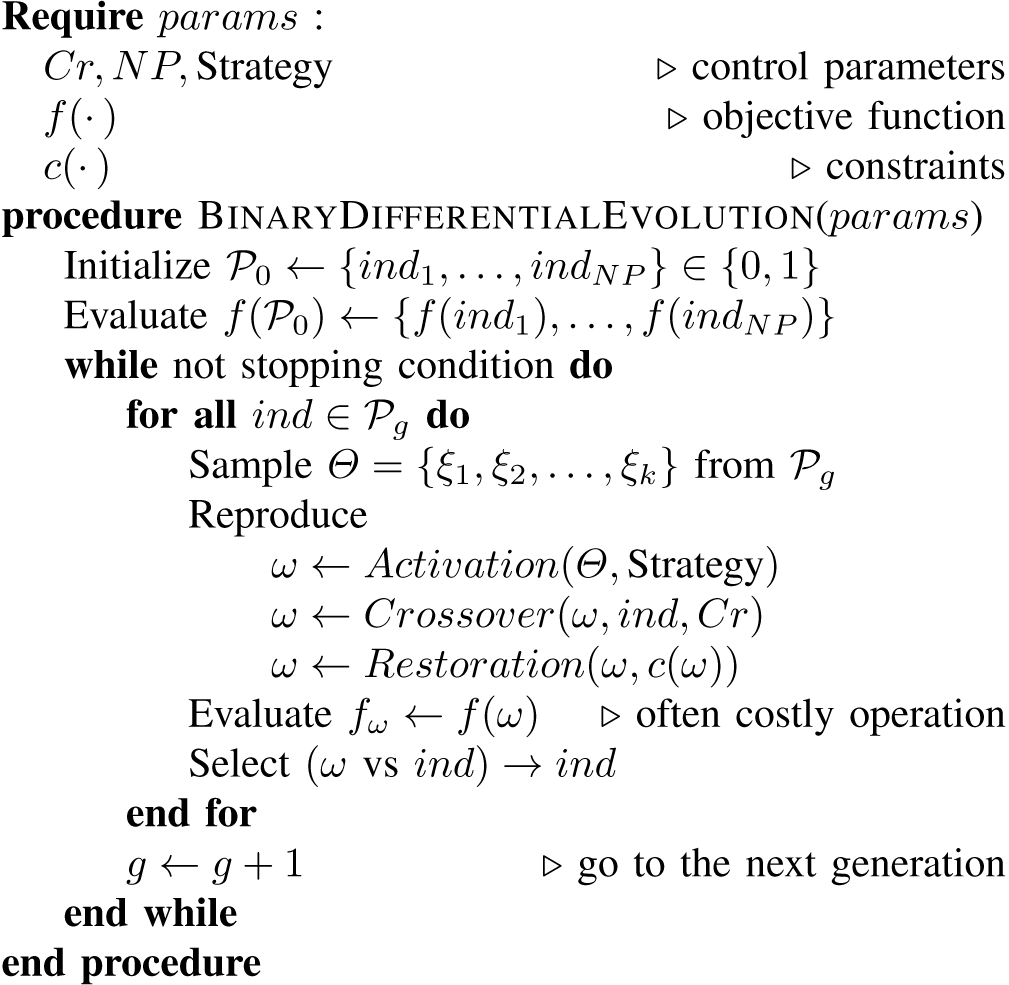

Here, we briefly describe the binary operators of reproducing cycle and exploring strategies designed for binary space.

#### a) Activation

This is the main operator for binary optimization we invented. It replaces the previous differentiation or also called differential mutation operator. As the previous one it is conform to the same formula [32]

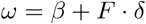

with base *β* vector and applied to it *δ* direction. But in a binary space we do not need the differentiation constant *F*, and the *δ* vector becomes an *activator α* which acts on the binary base vector *β*. Thus the binary formula becomes more simple

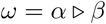

For any two sample from the population of solutions *ξ*_1_ and *ξ*_2_, the activator is computed as *α* = *ξ*_1_ – *ξ*_2_ and *α* ∈ {–1,0, + 1}^*n*^.

The activator *α* acts (▹) on *β* by switching on or off its *genes:*

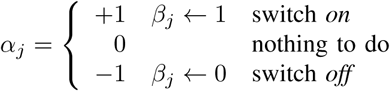

Thus, the activator controls the level of population diversity, and reduces proportionally when approaching the optimum.

#### b) Crossover

The role of crossover in this binary context is to preserve some parts of the individual to replace. Any conservative crossover can be used, for example

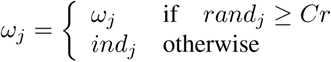

#### c) Restoration

After activation and crossover operations, very often, the constraints imposed on the *trial* individual *ω* are not respected. Many methods for handling constraints may be used to check the trial individual before evaluation. We show one of them, suitable for the budget constraint
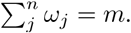

Let Δ = ∑_*j*_ *ω_j_* – *m* and Δ ≠ 0 then

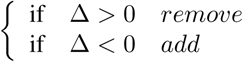

Δ of ones at random from/to the trial.

#### d) Strategies

For search space exploration, different techniques are available on how to choose and construct *α* and *β* from the set of sampled individuals *Θ*. Here we give four examples having different search properties.

1. <u>rand3</u> For each individual *ind* a set *Θ* of three other random individuals {*ξ*_1_ *ξ*_2_, *ξ*_3_} is sampled from the population 𝓟_*g*_. The activator *α* is defined as *α*;(*ξ*_1_, *ξ*_2_) = *ξ*_1_ – *ξ*_2_ and *β* = *ξ*_3_. So the trial

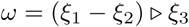 This strategy is good in most cases, especially for large-scale problems when an intensive exploration is needed.
2. <u>rand2best</u> Let *best* be the current best individual, that is *f* (*best*) ≤ *f*(*ind_i_*) ∀*ind_i_* ∈ 𝓟_*g*_. Two additional random individuals are extracted, *Θ* = {*ξ*_1_,*ξ*_2_}, to compute *α* = *ξ*_1_ – *ξ*_2_. *β* = *best*. Thus

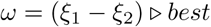 This strategy works well on mid-sized problems. It provides an excellent trade-off between exploration and exploitation.
3. <u>rand3dir</u> Three ordered individuals are sampled *Θ* = {*ξ*_1_, *ξ*_2_, *ξ*_3_}: *f* (*ξ*_1_) ≤ *f* (*ξ*_2_) ≤ *f* (*ξ*_3_). *α* = *ξ*_1_ – *ξ*_3_ and *β* = *ξ*_2_, so

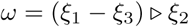 This strategy uses the gradient information of subspaces of the search space to control the exploration.
4. <u>rand2bestdir</u> The *best* individual is used in this strategy and there are two other randomly extracted individuals *Θ* = {*ξ*_1_, *ξ*_2_}, but this time the individuals are ordered *f*(*ξ*_1_) ≤ *f* (*ξ*_2_) and *best* ≠ *ξ*_1_. Then *α* = *ξ*_1_ – *ξ*_2_ and *β* = *best*, so

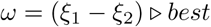 This strategy uses stochastic gradient information as well as the *best* individual. This is the combination of the previous two strategies in one. It has excellent convergence properties on small- and mid-sized problems.

### B. Optimization of hybrid testing

In hybrid breeding, it is usually not feasible to create and phenotype all possible hybrid combinations from a list of male and female inbreds. This problem can be reduced to the previous one by creating the virtual hybrid marker profile of all the hybrids of interest from their parents marker profile. Optimization is then done with the same approach as previously. One limit to investigate with such shortcut is the discrepancy between the optimization model which is a simplistic additive model and reality which should be closer to a model with distinct male and female alleles effect and an interaction component SCA (specific combining ability).

### C. Optimization of connected population design

Optimizing the design of a connected population is critical for recurrent selection scheme with genomic prediction.

A method is needed, using the potential parents marker profile, to evaluate the expected accuracy of a population before actually creating it. An optimization routine would use this evaluation method and optimize the design taking into account constraints on number of crosses, number of DH (doubled haploid) per crosses and total number of individuals. This is more difficult than the above described problems, because an efficient population design has to be identified before the population is actually created. Here we describe a solution to this problem combining analytical approaches and simulation of progeny. It goes as follow: for a given set of founders and their respective high density genotyping data, a very large number of DH progeny are simulated from all possible crosses from those founders. To evaluate a given design (which crosses and number of DH per cross), a number of populations are generated following that design. For each generated population the trace of the PEV is calculated using a fixed test set, made of a sample of each possible cross between founders. The objective function for the design is then the trace of *PEV* calculated on the fixed test set and averaged across generated populations. it can be further penalized by its standard error across generated populations to favor robust designs.

*Optimization algorithms:* The optimization problem is solved in two steps.

At the first step, a binary optimization method, similar to the one described in the previous Section II-A (see Alg. 1), is used to identify which pairs of founders to choose from all possible combinations respecting low *ϕ_lo_* and up *ϕ_up_* constraints on the number of selected pairs. There is an obvious limit to this first phase search: the optimization criteria increase monotonically with the population size. If the upper bound on number of crosses is equal to the total cross number, no cross will be eliminated in this first step. The largest number of crosses will then be favored.

At the second step, for each chosen pair from the first step, it is necessary to find the optimal number of DH to generate per cross (*ρ*), respecting technical boundary constraints *ρ* ∈ [*lo*, *up*] and total budget constraint ∑_*ρi*_ ≤ *total*.

For this purpose a new algorithm was created which is able to manipulate simultaneously mixed binary-integer variables. The integer variables are the *ρ* DH’ quantity by pair. The binary variables are switches that add or remove pairs of founders to the selection set depending on the actual number of DH.

We propose the alternative use of boundary constraints to incorporate binary optimization into the integer one. Let an integer variable^2^ *ρ* have its boundary constraints *ρ* ∈ [*lo*, *up*]. We introduce the notion of an *extended boundary* constraints [*lo*_2_,*up*] with *lo*_2_ = 2*lo* – *up*. This new extended constraint will act in three different modes:

1. <u>Boundary zone</u>: *neutral* integer optimization mode. If *ρ_i_* is in this range, nothing happens with its corresponding binary variable *b*_i_.
2. <u>Extended zone</u>: binary *exclusion.* If the *ρ_i_* value falls into the [*lo*_2_, *lo*) range the corresponding binary variable is switched off (*b_i_* ← 0). This variable is no more active.
3. <u>Out of zones</u>: binary *re*-*activation*. When *ρ_i_* takes values out of the extended range [*lo*_2_, *up*], the value of the corresponding binary variable is switched on, this variable becomes active again (*b_i_* ← 1).

The figure 1 resumes these modes.

**Fig. 1.**
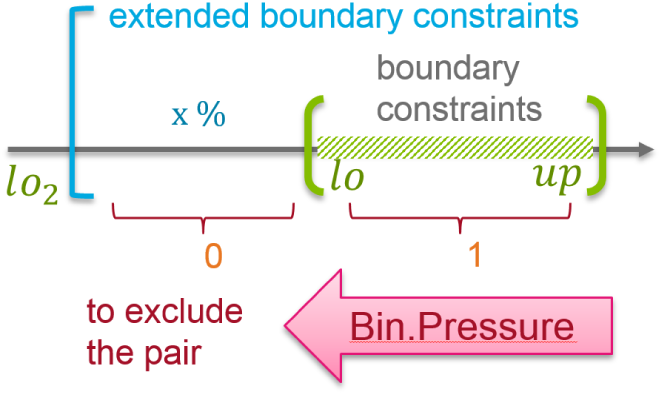
Extended boundary constraints.

The intensity of exclusion is called *binary pressure* (𝔅). The binary pressure can be controlled by varying the extended boundary position *lo*_2_ ← 𝔅 · *lo*_2_ with 𝔅 ∈ (0,1) or equivalently (0 – 100)% (see Fig. 2).

**Fig. 2.**
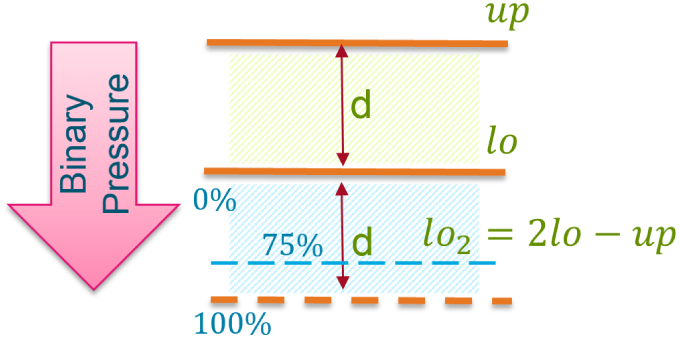
Binary pressure control.

When the binary variables are chosen it is necessary to respect the constraints for integer variables. Taking into account the need to effectively consume the total budget, the original methods of constraints handling inspired from sound engineering design is proposed. Let consider that the binary variables are a frequency spectrum and the integer variables are an amplitude at each frequency. *up* ~ 0dB clipping level. Thus the simulation of two principal sound engineering gears can be applied to correct the ”signal” *ρ* in its dynamic range (relative to constraints). Namely the *compressor* which will save the energy and the *amplifier* which will save the balance (see Fig. 3). Computing the ”space” *δ* = *total* – ∑*ρ_i_* with *ρ_i_* ∈ [*lo*, *up*] the following rules are applied

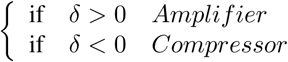

**Fig. 3.**
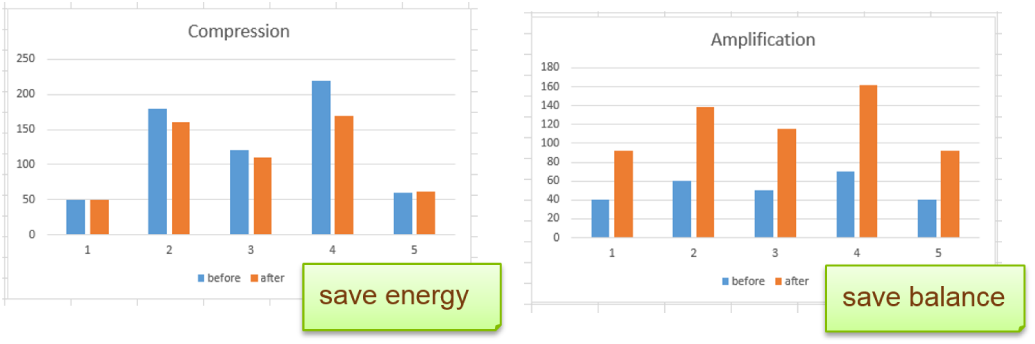
Phenotyping budget constraints handling.

### D. Validation with empirical data and simulations

To validate the described methods, real datasets were used and combined with simulations.

For selective phenotyping of existing genotypes, a dataset of 2522 elite winter wheat inbred lines were used, genotyped with more than 15000 SNPs (single nucleotide polymorphism) after quality control. 50, 100, 500 or 1000 individuals were selected using the method described earlier. Heritability for optimization was assumed to be 0.5. Solutions were generated using various number of eigenvectors. To validate the method, 200 markers were sampled as QTL. Corresponding effects were drawn from a normal distribution with a standard deviation of 1 and a true genetic value was computed for every individual from simulated QTL effects. Phenotype was generated from individual in calibration by adding to the true genetic value an error term drawn from a normal distribution. The error variance was adjusted to reach an heritability of 0.3, 0.5 or 0.8 to test the sensitivity of the optimization to the heritability assumption as in [20]. Using the generated phenotype, model of Eq. 1 was implemented using the R package rrBLUP [33], [34]. Markers sampled as QTL were excluded from the analysis. Accuracy is defined as the correlation between the predicted value for every individual and the simulated true genetic value for both observed and unobserved individuals. This procedure was repeated 100 times for optimized design of different sizes and random set of individuals of the same size.

For the hybrid design, a similar procedure was used. A dataset of 7147 real elite sunflower hybrids were used whose parents were genotyped with more than 10000 SNP markers after quality control. There was 631 males and 624 females. For validation, a true genetic value for each inbred and hybrid was generated using a genetic model closer to the expected genetic architecture.

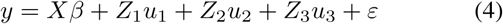

*y*, *β* and *ε* are as before. *u*_1_ and *u*_2_ are, respectively, male and female genetic effect such that
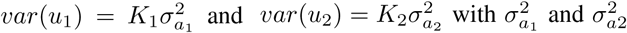
additive genetic variance in the male and female groups, respectively, and *K*_1_ and *K*_2_ relationship matrices, based on pedigree or markers for each group [9]. *u*_3_ is a dominance effect, such that
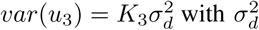
dominance variance and

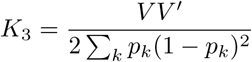

with *V* centered marker design matrix for the dominance effect such that each column of *V* corresponds to a marker with minor allele *a* of frequency *p* coded {*aa*, *Aa*, *AA*} = {–2*p*^2^, –2*p*(1 – *p*), –2*q*^2^} [35]. While *K*_1_ and *K*_2_ correspond to kinship between males and females for example, *K*_3_ has one row per hybrid. Heritability of the hybrid performance was simulated to be 0.5 but the fraction of variance due to dominance effect varied from 5% to 30% to test again the robustness of the optimization to the more likely true genetic architecture. 100 markers were sampled independently and taken as QTL for male additive effects, female additive effect and dominance effect. Simulations and analysis were implemented in R [36] using the package Asreml-R [37].

Last, for connected populations, six elite European flint lines were used in validation. 9273 SNPs passed basic quality control and were polymorphic among lines. Population design was optimized for a total budget of 600 DH, a maximum of 6 crosses with 50 to 300 DH per cross. 1000 solutions were tested in the first phase and 3000 in the second phase. The Oracle method was used in the first phase. For the second phase, differential evolution was used with the rand3 strategy. To validate the method, a simulation approach was used. 5, 10, 50, 100 or 1000 markers were used as QTL to simulate a true genetic value and test the robustness of the approach to the true genetic architecture. Heritability was set to 0.5. Simulations and analysis were implemented in R [36]. The generated phenotype was analyzed using (Eq. 3) implemented with the package rrBLUP [33], [34]. The optimized design was compared to classical designs depicted in figure 4 and to a set of 100 design generated at random with a total population size of 600 DH. Efficiency of each design was measured using the following measures:

**Fig. 4.**
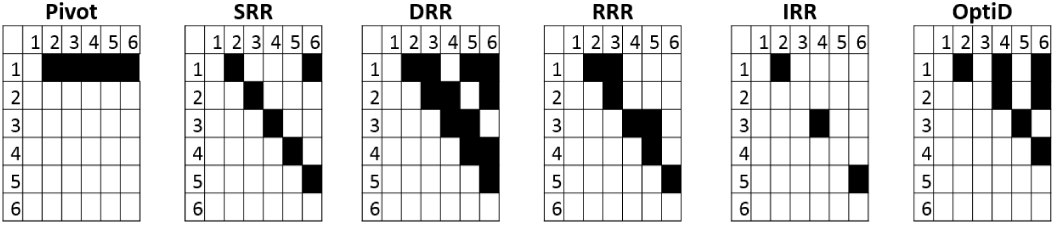
Mating designs considered in this study: Pivot, single round robin (SRR), double round-robin (DRR), reduced round-robin (RRR), independent round-robin (IRR), Optimized. Black squares indicate the crosses from which DH are derived.

- Accuracy of prediction of the true breeding value for the generated population (also phenotyped).
- Accuracy of prediction of the true breeding value of an independent unobserved set of 100 DH from a single randomly chosen cross.
- Accuracy of prediction of the true breeding value of an independent unobserved set made of 10 DH from every possible crosses between the possible founders.
- Root mean square of error (RMSE) between the estimated alleles effects and the true simulated effect when only the simulated causal markers are used in the analysis. In each design, non segregating QTL were either discarded from the computation of the RMSE of their estimated effect was considered to be 0. This was only computed in simulations with at least 50 QTLs.

Correlation between the efficiency measures obtained with the different design and the objective function were also analyzed. All optimization routine were implemented in Fortran and multi-threaded

## III. RESULTS

### A. Optimization of selective phenotyping

Optimisation took 4.5 hours with 25 eigenvectors to select a subset of 100 individuals from 2522. 3 millions solutions were evaluated by the algorithm. For computation, only four cores (threads) of a Intel Xeon E5-4627 v3 processors operating under Windows Server 2012 R2 were used. The figure 5 presents the convergence rate as a function of the number of evaluations for the first 600 000 evaluations. No further improved solution were found in the next 2.4 millions evaluated solutions.

**Fig. 5.**
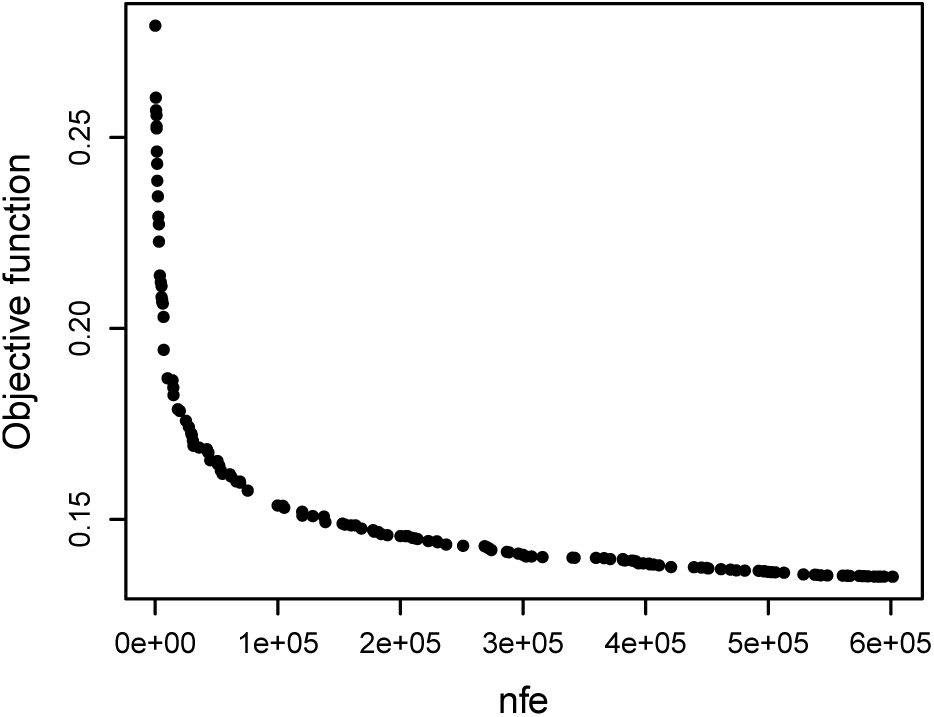
Convergence rate: *nfe* - number of function evaluation for the first 600 000 evaluations to select 100 individuals from 2522 from the wheat dataset

Validation showed, for a given subset size, that using a large number of eigenvectors was detrimental to the quality of the solution and too few eigenvectors gave results worse than random solutions (figure 6). After tests on a number of datasets, using a fourth of the subset size as number of eigenvectors seems to be a reliable heuristic. In the rest of the paper, results are presented using this rule of thumb.

**Fig. 6.**
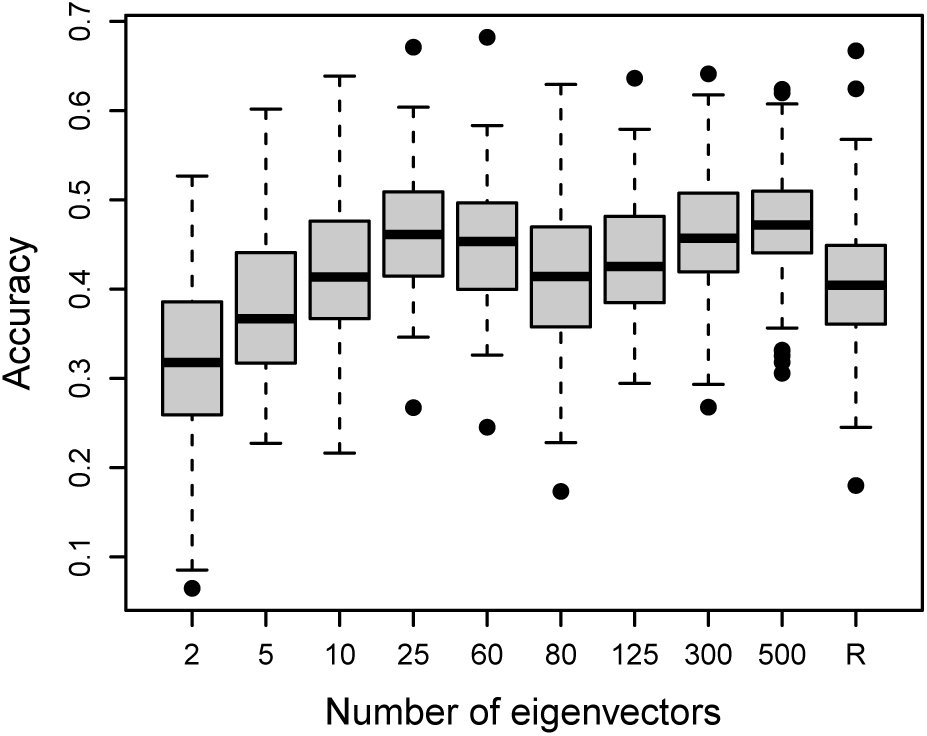
Comparison of accuracy obtained with varying number of eigenvectors and random solution (R), for a subset size of 100, heritability of 0.5 in optimization and simulations, for the wheat dataset.

For small subset of individuals compared to the total population size (figure 7), the optimized set gave a significantly higher accuracy than random set of individuals. Accuracy gain reached 20% when 50 individuals were selected from 2522 and dropped to 2% when 500 individuals were selected. Similar results were obtained when trait heritability in simulation was lower (0.3) or higher (0.8) than assumed in optimization (0.5) as already reported in [20].

**Fig. 7.**
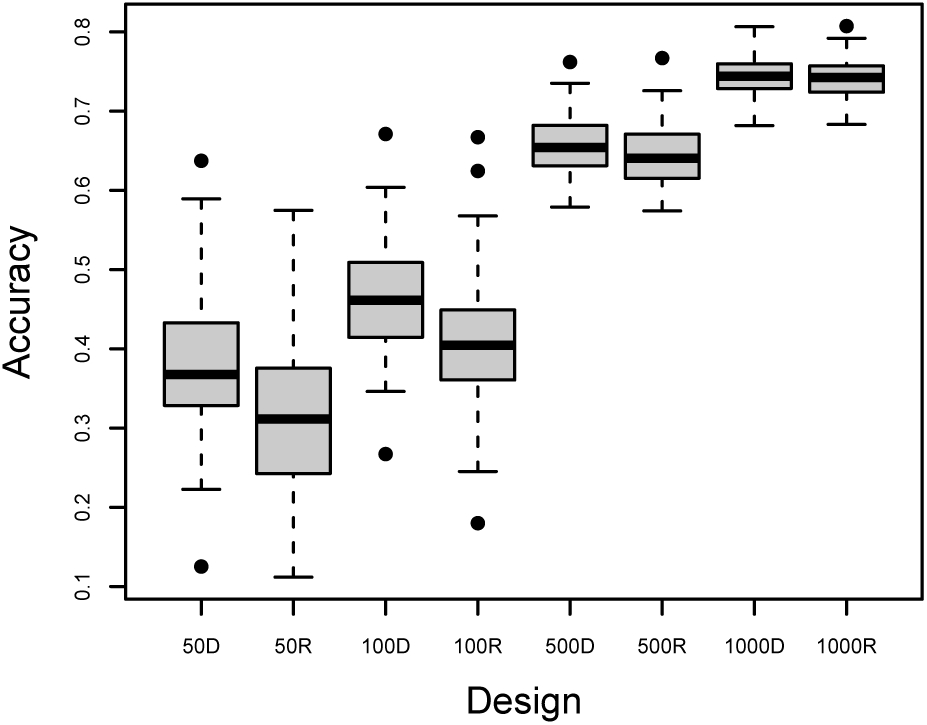
Comparison of accuracy obtained with random sets (50R, 100R, 500R, 1000R) or optimized set of different sizes (50D, 100D, 500D, 1000D) heritability of 0.5 in optimization and simulations for the wheat dataset

### B. Optimization of hybrid testing

Similarly when the same algorithm was used to select a subset of sunflower hybrids to phenotype, the optimized set gave a significantly higher accuracy than random set of individuals when the validation model was assuming a perse genetic architecture (figure 8). Accuracy gains reached 21% when 50 hybrids were selected from 7147 and dropped to less than 1% when 3000 hybrids were selected.

**Fig. 8.**
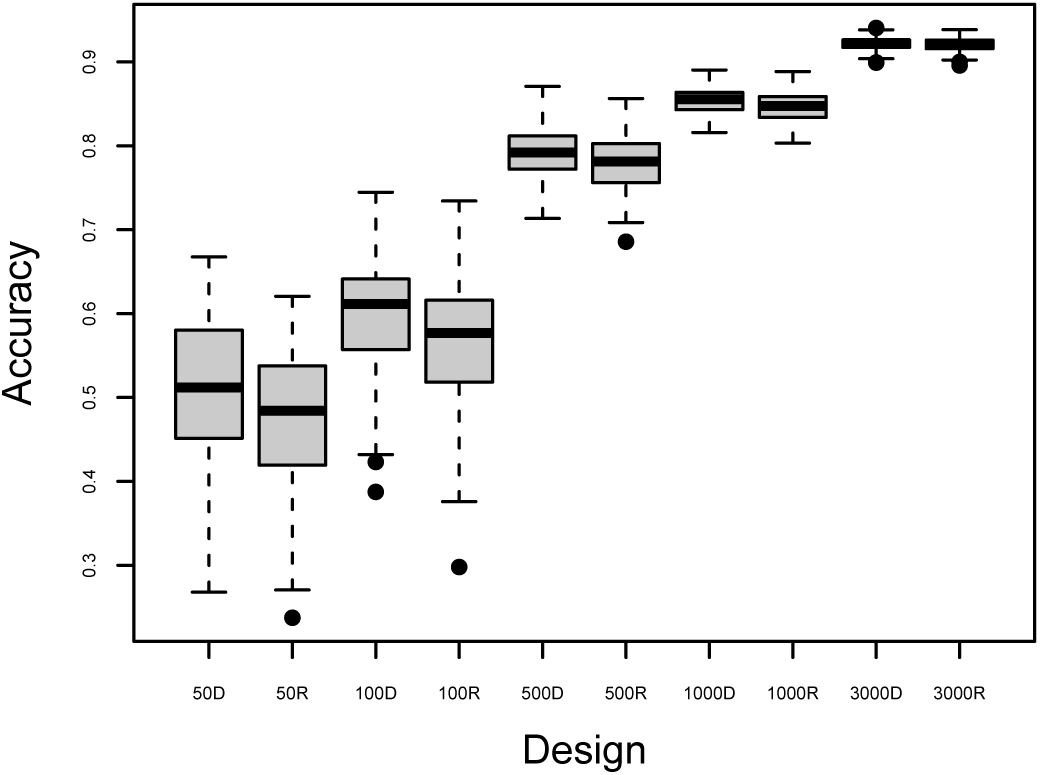
Comparison of accuracy obtained with random sets (50R, 100R, 500R, 1000R, 3000R) or optimized set of different sizes (50D, 100D, 500D, 1000R, 3000R) heritability of 0.5 in optimization and simulations for the sunflower hybrid dataset and a perse model for validation

If a more realistic model was used to simulate true genetic effects and analyze the simulated data, accuracy gains were still observed (figure 9). Accuracy gains reached 13% when 50 hybrids were selected from 7147 and dropped to less than 1% when 3000 hybrids were selected. Similar accuracy gains were also observed for each genetic effect (males, females and dominance effect).

**Fig. 9.**
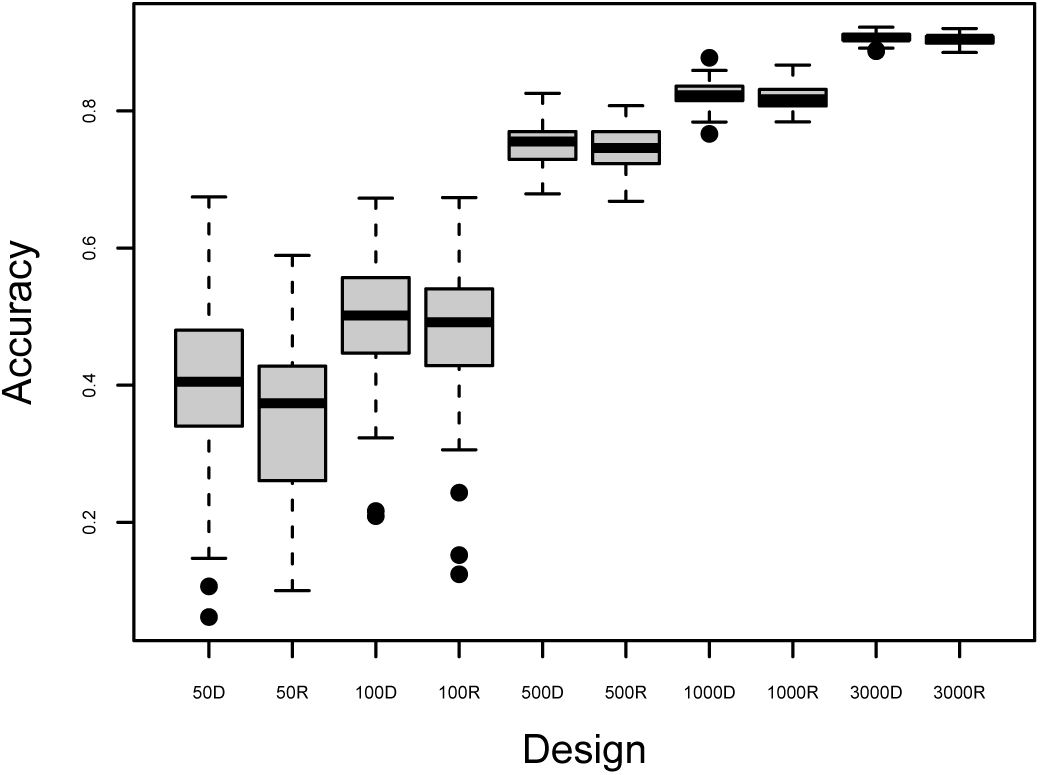
. Comparison of accuracy obtained with random sets (50R, 100R, 500R, 1000R, 3000R) or optimized set of different sizes (50D, 100D, 500D, 1000R, 3000R) heritability of 0.5 in optimization and simulations for the sunflower hybrid dataset, independent males and females genetic effects and a 20% dominance variance

### C. Optimization of connected population design

Optimization took 6.1 hours with 25 eigenvectors to identify a design with 600 DH in total, from 1 to 10 crosses with a possible cross size between 50 and 600. Each solution was evaluated by the algorithm using 100 replicates. 1000 solutions were evaluated in the first step and 3000 in the second step. For computation, only six cores (threads) of a Intel Xeon E5-4627 v3 processors operating under Windows Server 2012 R2 were used. The identified optimal design is shown in figure 4. The figure 10 presents the convergence rate as a function of the number of evaluations. The vertical line indicates the start of the second step.

**Fig. 10.**
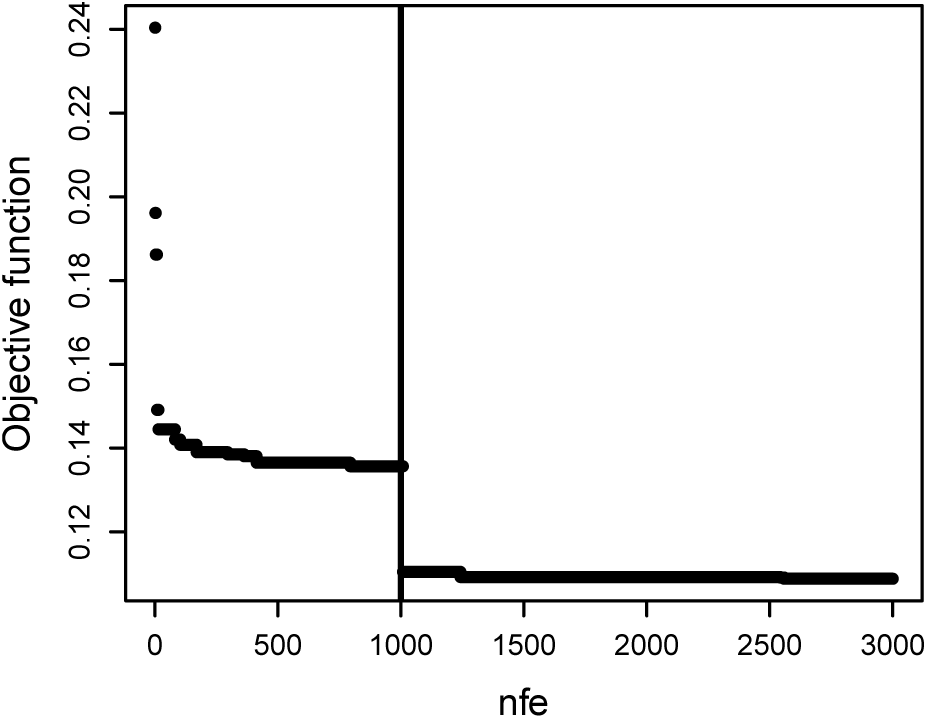
Convergence rate: *nfe* - number of function evaluations. The vertical line indicates the start of the second step.

The predictive power of the objective function can be assessed on figure 11. Each dot on the figures is a population design, all have the same total population size of 600 DH.

**Fig. 11.**
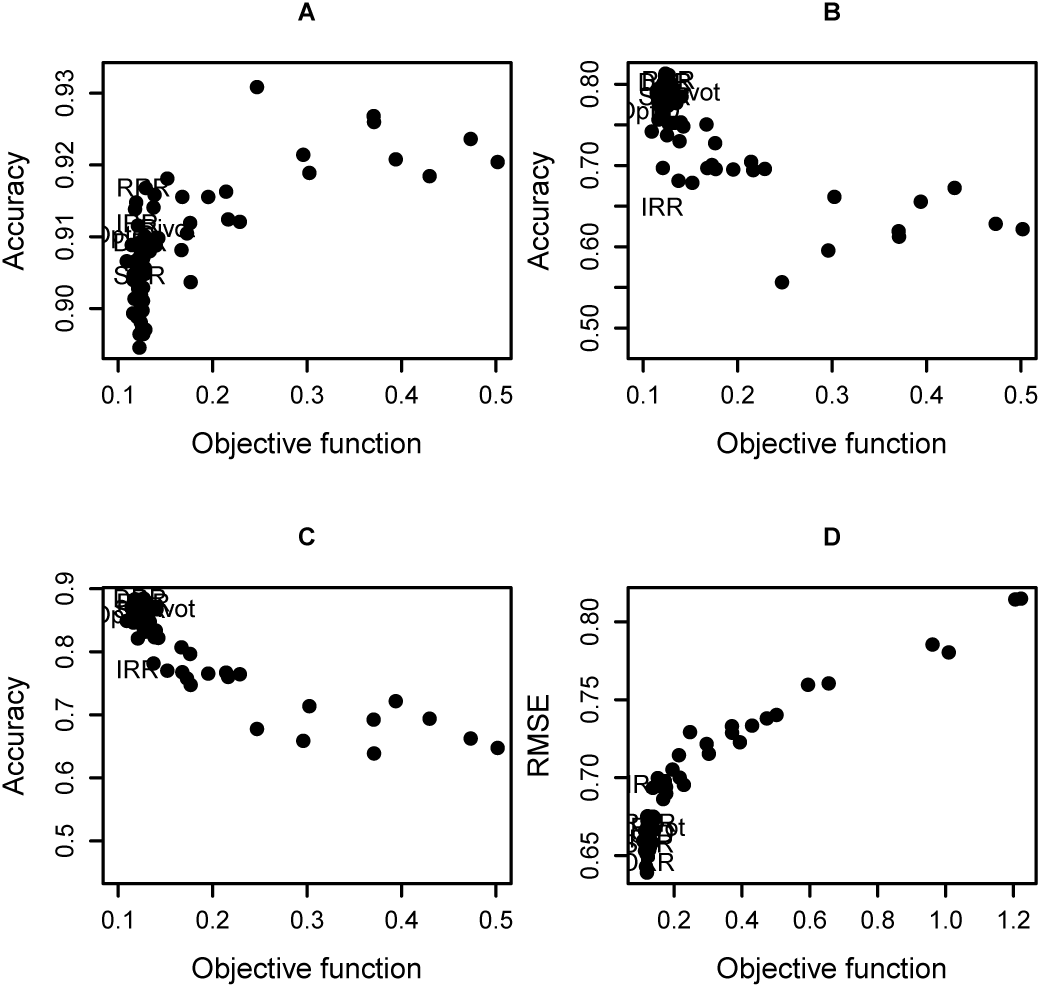
Design performance metrics as a function of the objective function for classical design, optimized mating design: Pivot, single round robin (SRR), double round-robin (DRR), reduced round-robin (RRR), independent round-robin (IRR), Optimized and randomly generated design, all with the same total population size of 600 DH. **A** Accuracy of prediction of the true breeding value for the generated population (also phenotyped). **B** Accuracy of prediction of the true breeding value of an independent unobserved set of 100 DH from a single randomly chosen cross. **C** Accuracy of prediction of the true breeding value of an independent unobserved set made of 10 DH from every possible crosses between the parents. **D** Root mean square of error (RMSE) between the estimated alleles effects and the true simulated effect when only the simulated causal markers are used in the analysis. The effect of non segregating QTL was considered to be 0. Results are averaged across simulations with different number of QTL: 5, 10, 50, 100 or 1000, each repeated 100 times.

The objective function is well correlated with three of the four metrics: Accuracy of prediction of the true breeding value of an independent unobserved set of 100 DH from a single randomly chosen cross, accuracy of prediction of the true breeding value of an independent unobserved set made of 10 DH from every possible crosses between the parents and with the root mean square of error (RMSE) between the estimated alleles effects and the true simulated effect when only the simulated causal markers are used in the analysis. However, the objective function is negatively correlated with the accuracy of prediction of the true breeding value for the generated population which is also phenotyped. But the range of accuracies was small for this specific metric (0.89 to 0.93). Populations with a large objective function had fewer crosses and were less connected.

It is also clear from figure 11 that a small difference in the objective function has a limited impact on the performance of the design. This is confirmed by a more in-depth comparison of the classical and optimized design presented on figure 12. Differences on the various performance metrics were very small except for the IRR design which is also the most disconnected one. The objective function differences between design were also very small ranging from 0.105 for the optimized design, 0.126 for the IRR design to 0.15 for the pivot design.

**Fig. 12.**
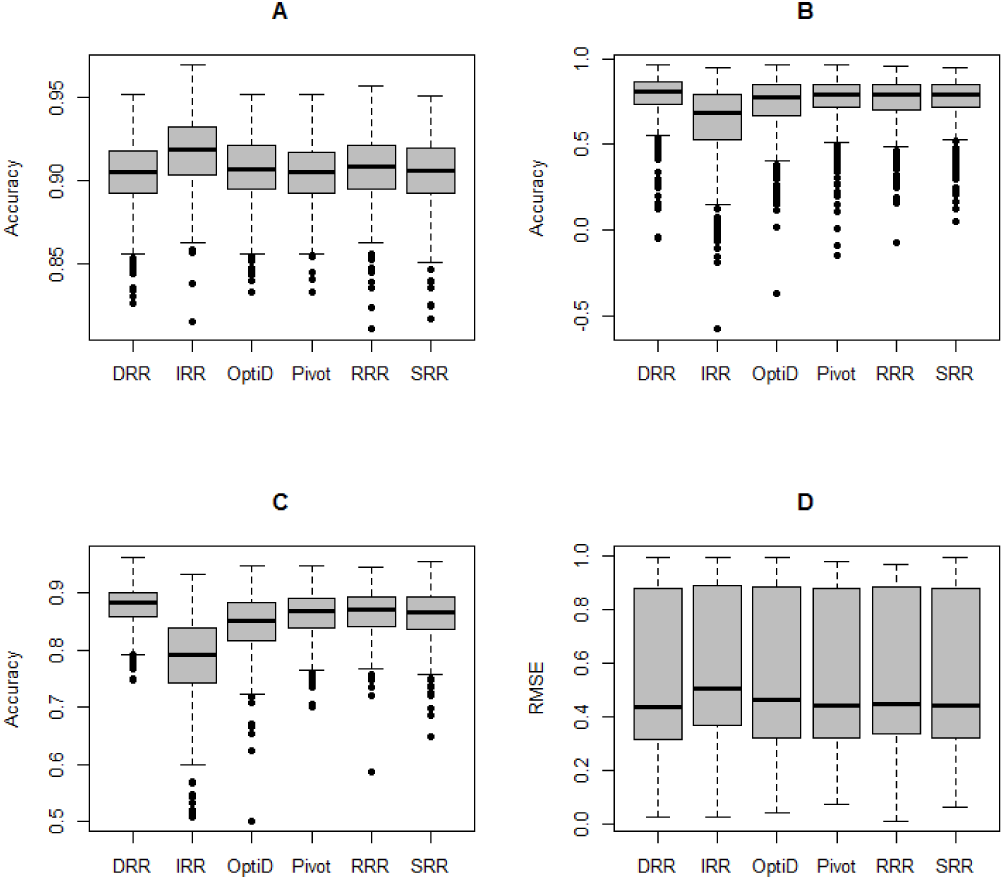
Comparison of classical design and optimized mating design: Pivot, single round robin (SRR), double round-robin (DRR), reduced round-robin (RRR), independent round-robin (IRR), Optimized. **A** Accuracy of prediction of the true breeding value for the generated population (also phenotyped). **B** Accuracy of prediction of the true breeding value of an independent unobserved set of 100 DH from a single randomly chosen cross. **C** Accuracy of prediction of the true breeding value of an independent unobserved set made of 10 DH from every possible crosses between the parents. **D** Root mean square of error (RMSE) between the estimated alleles effects and the true simulated effect when only the simulated causal markers are used in the analysis. The effect of non segregating QTL was considered to be 0. Results are presented together for the different number of QTL: 5, 10, 50, 100 or 1000, each repeated 100 times.

If instead of comparing design with the same total population size, we consider a set of 100 randomly generated design with a population size between 100 and 1000 DH, figure 13, the same conclusion are valid. With a difference for the accuracy of prediction of the true breeding value for the generated population. Now, the objective function is also positively correlated with this metrics.

**Fig. 13.**
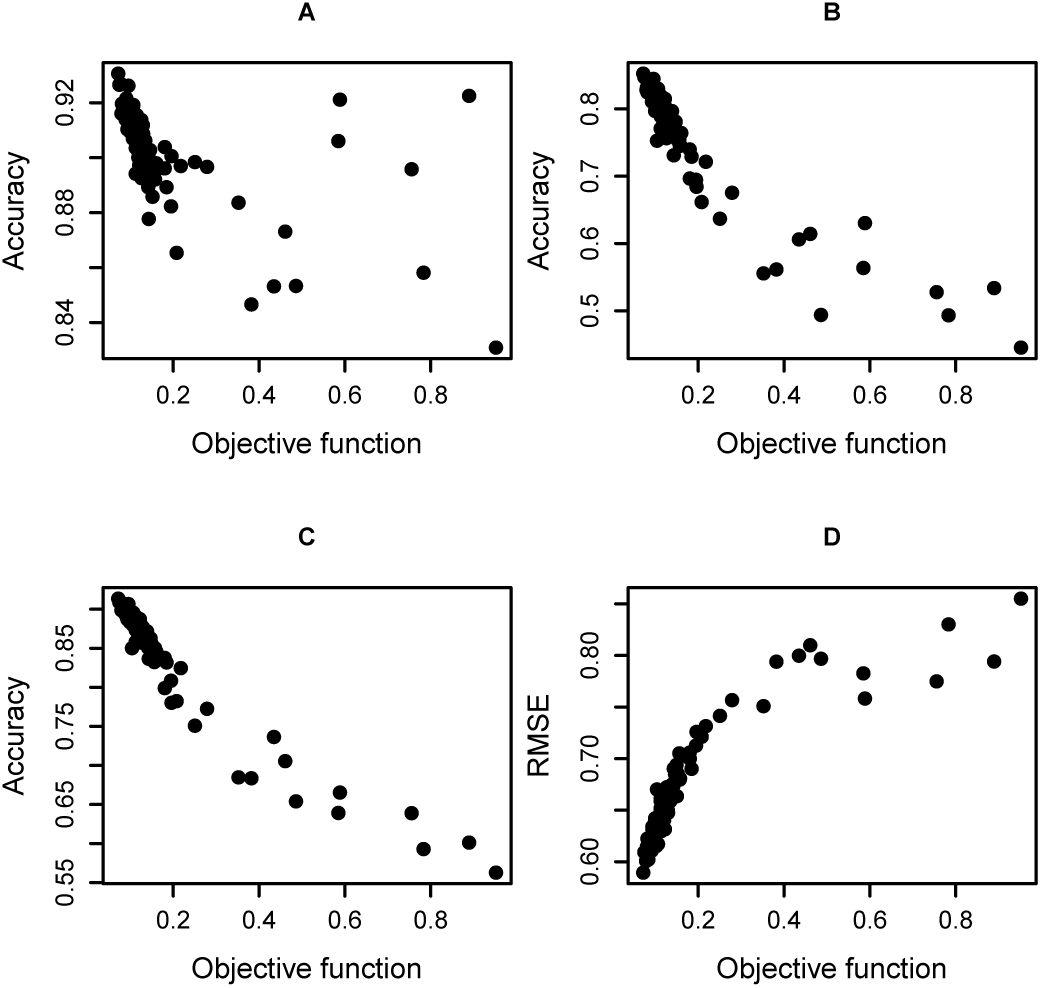
Design performance metrics as a function of the objective for randomly generated design with a population size varying between 100 and 1000 DH. **A** Accuracy of prediction of the true breeding value for the generated population (also phenotyped). **B** Accuracy of prediction of the true breeding value of an independent unobserved set of 100 DH from a single randomly chosen cross. **C** Accuracy of prediction of the true breeding value of an independent unobserved set made of 10 DH from every possible crosses between the parents. **D** Root mean square of error (RMSE) between the estimated alleles effects and the true simulated effect when only the simulated causal markers are used in the analysis. The effect of non segregating QTL was considered to be 0. Results are averaged across simulations with different number of QTL: 5, 10, 50, 100 or 1000, each repeated 100 times.

Those results were also confirmed on a number of other populations.

## IV. DISCUSSION

### A. Optimization of selective phenotyping and hybrid design

Algorithms for selective phenotyping were proposed by [20]. The method presented here is very similar but use modified differential evolution instead of simple interchanges to identify an optimal subset of individuals to phenotype. [21] combined the approach of [20] with stratified sampling to improve the results in structured populations. [22] used an uniform sampling approach for the same objective and compared those methods along with a few simple others. Contrary to our results, differences between methods were smaller at small training set sizes than at larger training set sizes. Our opposite results are probably due to a superior optimization algorithm. The optimization problem being more difficult for a small rather than a large training set size. [22] reported that uniform sampling method also provided more robust results, probably also a consequence of the optimization routine used. If the optimization method is not very effective, using a simpler criteria ensuring uniform coverage of the genetic space has more chances of providing a reliable training set. [28] on the contrary used a genetic algorithm for optimization and reported larger accuracy gains for smaller populations. In [20] and [28] much fewer iterations of the optimization algorithms were used. [28] reported no improvements after 200 evaluations and [20] mostly use repeats of a simple exchange algorithm to explore the search space. With the algorithm presented here, better solutions were still found up to 600 000 solution evaluations (figure 5). This suggests that our optimization procedure is superior. Algorithm was also further developed to handle constraints (individuals that have to be observed or individuals to exclude from selection but with interest in prediction).

Quite surprisingly, despite a simplistic optimization model, assuming additive genetic effects, the algorithm provided good results to identify a subset of hybrids to phenotype. This despite a more realistic genetic architecture used in validation, including independent males and females additive effects and a dominance effect. Obviously results could be further improved here by deriving a more realistic optimization criteria for an hybrid genetic model. The limited impact of the true genetic architecture on the optimization suggests that this type of approach should be considered with pragmatism in the same way that model based design is used to generate reasonable and efficient field trial designs [25].

The number of eigenvectors used to approximate the objective function had a large impact on the efficiency of optimization. With few eigenvectors, results were better for selection of small subsets than with many eigenvectors, and significantly worse than random for large subsets. Taking a fourth of the subset size as number of eigenvectors seems to be a robust heuristic. Using few eigenvectors smooths the objective function and accelerates convergence but might oversimplify it. With a small subset, this simplification does not have a penalty because the subset size limits the sampling of the genetic space. In addition the optimization problem is more difficult for a small subset due to a much larger search space. There is then an advantage to simplify the objective function to ease convergence. On the contrary for a large subset size, the genetic space can be better sampled. Using few eigenvectors can oversimplify the objective function and lead to a biased sample, worse than random sampling.

### B. Optimization of connected population design

In the context of QTL mapping a number of studies focused on identifying efficient connected population design for QTL detection. [18] used simulation and compared a number of design including full diallel, factorial (reduced diallel) NAM (nested association mapping), called pivot in this study, round robin, double round robin, reduced round robin and so-called distance based design with selection of a set of crosses maximizing genetic dissimilarity. The least powerful design in his simulations were NAM and distance based ones. The most powerful were diallel, factorial and double round robin. Here, in the context of genomic prediction we broadly confirmed those results and our optimization criteria is able to correctly rank population design for suitability for genomic prediction. Maximizing accuracy of marker effects estimates will improve the efficiency of recurrent selection with genomic prediction from those optimized populations. The method presented here, can then be used to optimize a population design, taking into account practical constraints, but also to compare alternative population designs with varying number of crosses, connectivity and total size. This makes it a decision support tool for population design for genomic prediction and probably QTL detection. Basic questions such as how much accuracy will be lost by removing a given cross from the design or by reducing the total population size can be easily answered. The results also show that a small difference in objective function between high quality design (small objective function) has a limited impact on design quality. This is a critical point in comparing design alternative. We recommend to systematically compare the optimized design with classical alternatives to have a reference point.

Results of figure 11, points to a fundamental contradiction to overcome in designing population for genomic prediction: for a given population size, a design with very few crosses (even down to a single cross) will deliver the maximum accuracy for the generated population. But it will be very inefficient to predict unobserved individuals from other crosses or accurately estimate the effects of the QTL segregating between the possible parents of the population. The proposed objective function is addressing the second point. It is thus not surprising that the identified optimal design or classical design will be slightly less efficient in accuracy for the generated population but much more accurate to predict unobserved individuals from other unobserved crosses or to estimate the effects of the QTL segregating in the possible parents.

## Footnotes

1 Do not confuse algorithm’s terms with previous genetic population and individuals to be phenotyped. Historically the terms are the same but their meanings are very different.

2 Continuous variables can be enlarged to mixed binary-continuous optimization by the same way.

